# Encoding of environmental cues in central amygdala neurons during foraging

**DOI:** 10.1101/2020.09.28.313056

**Authors:** Marion Ponserre, Federica Fermani, Rüdiger Klein

**Author notes:** Correspondence (R.K.). These authors contributed equally to this work (Marion Ponserre, Federica Fermani).

## Abstract

In order to successfully forage in an environment filled with rewards and threats, animals need to rely on familiar structures of their environment that signal food availability. The central amygdala (CeA) is known to mediate a panoply of consummatory and defensive behaviors, yet how specific activity patterns within CeA subpopulations guide optimal choices is incompletely understood. In a paradigm of appetitive conditioning in which mice freely forage for food across a continuum of cues, we find that two major subpopulations of CeA neurons, Somatostatin-positive (CeA^Sst^) and protein kinase Cδ-positive (CeA^PKCδ^) neurons can assign motivational properties to environmental cues and encode memory of goal location. While the proportion of food responsive cells was higher within CeA^Sst^ than CeA^PKCδ^ neurons, only the activities of CeA^PKCδ^, but not CeA^Sst^, neurons were required for learning of contextual food cues. Since CeA^PKCδ^ neurons are known to promote a range of defensive behaviors, our findings point to a model in which CeA circuit components are not organized in specialized functional units but can process both aversive and rewarding information in a context and experience dependent manner.

**HIGHLIGHTS:** Two populations of central amygdala (CeA) neurons, CeA^PKCδ^ and CeA^Sst^ neurons can assign motivational properties to environmental cues and encode memory of goal location.

The proportion of food responsive cells was higher among CeA^Sst^, than CeA^PKCδ^ neurons.

The activities of CeA^PKCδ^, but not CeA^Sst^, neurons are required for learning of contextual food cues. CeA^PKCδ^ neurons represent a “general encoding” population selecting defensive and appetitive responses depending on context.

## INTRODUCTION

An organism’s survival depends heavily on its ability to evaluate whether environmental cues predict a threat or an opportunity. The CeA is important for this competence and contains numerous genetically distinct subpopulations of neurons that regulate approach and avoidance responses to positively and negatively valenced stimuli (Ehrlich et al., 2009; Fadok et al., 2018; Herry and Johansen, 2014; Janak and Tye, 2015; Pare and Duvarci, 2012). Three scenarios seem possible: First, the CeA contains “general encoding” populations for a variety of stimuli, and, depending on context, they will select the appropriate circuit elements to generate a defensive or approach response. Second, the CeA contains broad populations that relate to stimuli of either positive or negative valence. For instance, a “broad aversive” population relates to painful and threatening, but not rewarding stimuli. Third, the CeA contains distinct populations that promote specifically one class of behavior. For example, a “distinct appetitive” population mediates approach responses to food, but not other rewarding stimuli.

There is currently support for all three models. CeA^Sst^ neurons may represent a “general encoding” population, relating to appetitive stimuli (Kim et al., 2017; Wilson et al., 2019) and contributing to the generation of defensive behaviors (Li et al., 2013; Penzo et al., 2015; Yu et al., 2016). CeA^PKCδ^ neurons could be characterized as “broad aversive”, since these neurons promote aversive and suppress appetitive responses (Cai et al., 2014; Haubensak et al., 2010; Kim et al., 2017; Wilson et al., 2019; Yu et al., 2017). CeA neurons marked by expression of prepronociceptin represent “distinct appetitive” cells, mediating food intake, but not liquids (Hardaway et al., 2019).

To provide further evidence for these models and understand how the surrounding environment influences CeA neuronal activity to drive behavioral selection, we probed the function and activity patterns of CeA^PKCδ^ and CeA^Sst^ neurons during a foraging task. The CeA has been previously shown to acquire responses to signals of food delivery, but the contribution of these two CeA subpopulations has remained unclear (Corbit and Balleine, 2005; Gallagher et al., 1990; Han et al., 1997, 1999; Lee et al., 2005; McDannald et al., 2004). Especially how these appetitive memories are represented in a semi-naturalistic setting where the animal freely navigates and is forced to engage goal-oriented actions is unknown.

Here, we found that CeA^PKCδ^ neurons, previously related to aversive behavior, were required for approach behavior to a context that signals food availability, whereas CeA^Sst^ neurons were not. We performed Ca^2+^ imaging to evaluate neuronal activity changes when the mice freely decided to switch between environments. We observed that after learning, a subset of these two populations had become memory neurons that had developed specific activities to the positive context. Our findings suggest a model in which CeA^Sst^ neuron activity encodes information directly related to food consumption, whereas CeA^PKCδ^ cells store memories about the salience of the food reward. These findings support the first model in which PKCδ+ neurons represent a “general encoding” population selecting defensive and appetitive responses depending on circumstances and shifts our focus away from models of CeA function in which subpopulations represent hardwired and specialized functional units.

## RESULTS

### Inhibition of CeA^PKCδ^ neurons impairs contextual appetitive conditioning

To assess the contributions of CeA subpopulations to the acquisition of contextual appetitive memories, we developed a place preference paradigm inspired by previous work (Stern et al., 2018). On the first day (Figure 1A), *ad libitum* fed mice were habituated to an arena with two-chambers that differ in many features. We then performed four training sessions in which the mice were food restricted and sequestered in each chamber for an equal amount of time, but were given access to food only in the chamber that was the least preferred during habituation (positive context). On the following day, food was removed from the positive context and mice were allowed to freely navigate between the two chambers. We measured the amount of time the mice spent in the positive context as a readout for contextual appetitive conditioning. Typically, WT mice learned the task well, spending on average 71.9 +/-7.2 % of their time in the positive context (Figure 1B).

**Figure 1:**
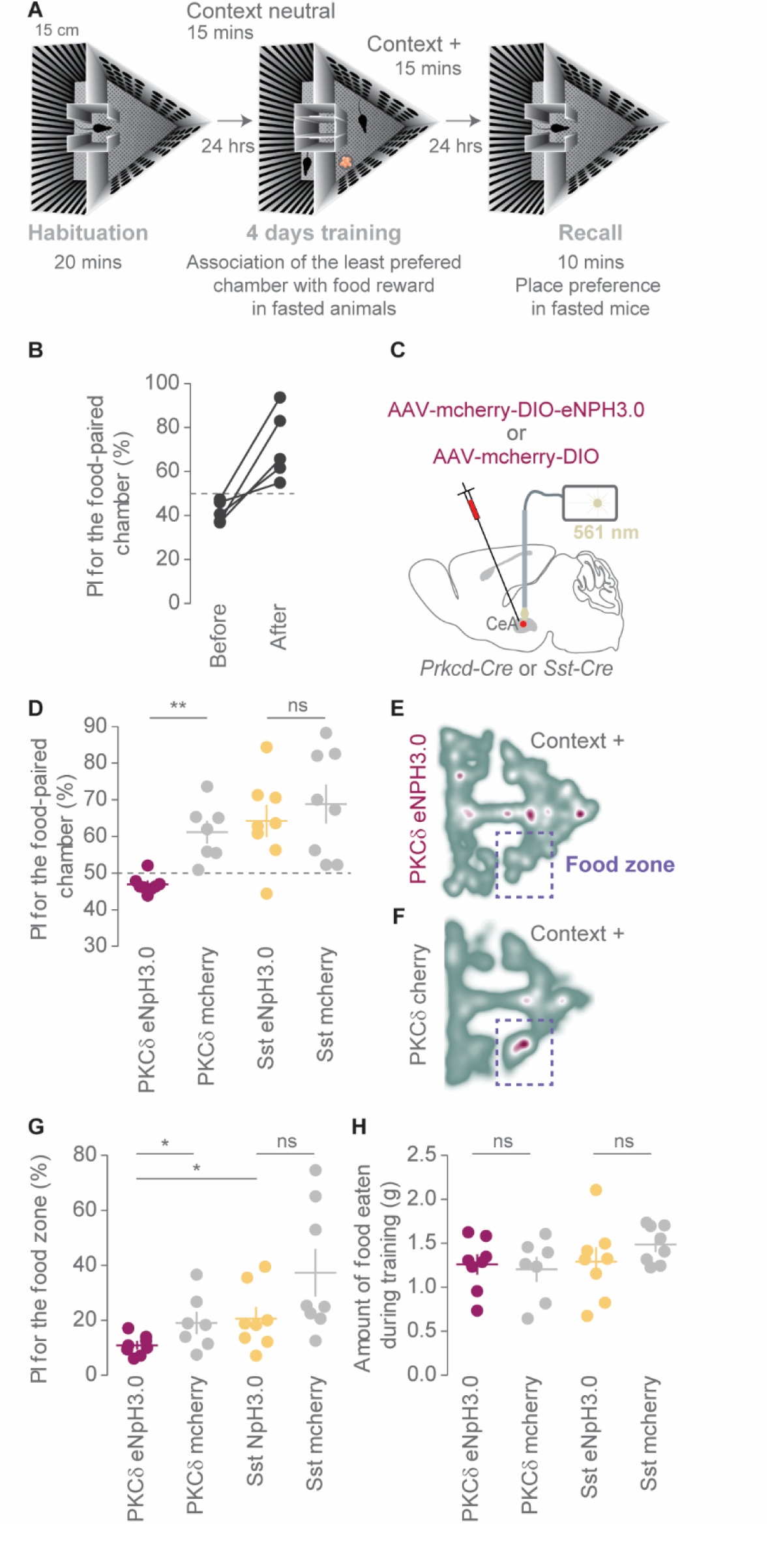
Inhibition of CeA^PKCδ^ neurons impairs contextual appetitive conditioning. (A) Place preference behavioral paradigm. (B) Preference index (PI) for the context+ before and after conditioning for WT animals (n= 5 mice). (C) Optogenetic inhibition of CeA neurons. The CeA of Prkcd-Cre or Sst-Cre mice was transduced with a Cre-dependent AAV expressing eNPH3.0-mcherry or mcherry alone. During each training session, CeA neurons were photostimulated *in vivo* with constant yellow light. (D) PI for the food-paired chamber (context+) during recall (Mann-Withney U test, for PKCδ group comparison: U = 1, P = 0.0021 and for Sst group comparison: U = 29, P = 0.7927). (E-F) Representative heat maps of the behavior of individual PKCδ-eNpH3.0 and PKCδ-mcherry mice during recall. Green represents the minimum and purple the maximum per-pixel frequency. (G) PI for the food zone during recall (Mann-Withney U test, for PKCδ group comparison: U = 10.5, P = 0.0489 and for Sst group comparison: U = 14, P = 0.0650). (H) Cumulative amount of food eaten during the four training days (two-tailed unpaired t-test, for PKCδ group comparison: t(13) = 0.3398, P = 0.7394 and for Sst group comparison: t(14) =1.141, P = 0.2730). (n = 8 PKCδ-eNpH3.0 and 7 PKCδ-mcherry mice and n = 8 Sst-eNpH3.0 and n = 8 Sst-mcherry mice). Bar graphs show mean +/- s.e.m and each dot is the quantification of a single animal; ns, not significant, *P < 0.05, **P < 0. 01.

To optogenetically inhibit either CeA^PKCδ^ or CeA^Sst^ neurons we infected the CeAs of *Prkcd-Cre* or *Sst-Cre* animals with Cre-dependent halorhodopsin-expressing virus (eNpHR3.0-mCherry) and placed optic fibers bilaterally above the CeA (Figure 1C and Figures S1A-B). Control mice expressing mCherry behaved as WT mice although their preference index after learning was slightly lower (61.4 +/-2.9 % for PKCδ mcherry and 69 +/-5.1 % for Sst mcherry mice) (Figures 1D and S1C-D). Silencing of CeA^Sst^ neurons did not alter the behavior (Figures 1D and S1C). In contrast, CeA^PKCδ^ neuron-inhibited mice showed poor learning performance, lacking a preference for the positive context during recall (Figure 1D and S1D). In addition, CeA^PKCδ^ neuron-inhibited animals spent significantly less time in the food zone of the positive context (Figure 1E-G), even though they visited it with the same frequency as control mice (Figure S1E). This is in opposition to CeA^Sst^ neuron-inhibited mice, which showed similar numbers of visits and preference for the food zone as controls (Figure 1G and S1E).

Several other parameters that may have influenced the outcome of this learning paradigm were found to be similar between the experimental groups, including the amount of food eaten during training (Figure 1H), velocity during recall (Figure S1F), and the degree of anxiety as measured by duration and entries to the center, and velocity in an open field task (Figures S1G-M). Overall, these results suggest that CeA^PKCδ^ neurons may play a role in forming associations between contextual cues and food availability.

### *In vivo* recordings of CeA^PKCδ^ and CeA^Sst^ neuron activity during appetitive conditioning

To understand how CeA^PKCδ^ and CeA^Sst^ neuron activities differentially contribute to this behavior, we tracked the Ca^2+^ dynamics of these two populations during all phases of the task. We injected the CeA of Prkcd-Cre and Sst-Cre mice with an adeno-associated virus (AAV) expressing GCaMP6s in a Cre-dependent manner and recorded the dynamics of GCaMP6s fluorescent signals using a miniature microscope (Figure 2A-B and Figure S2A-B). Mice were subjected to the same behavioral task and at the end of the recall, we added a food pellet in the positive context to make sure that we could functionally tag neurons that responded to food intake.

**Figure 2:**
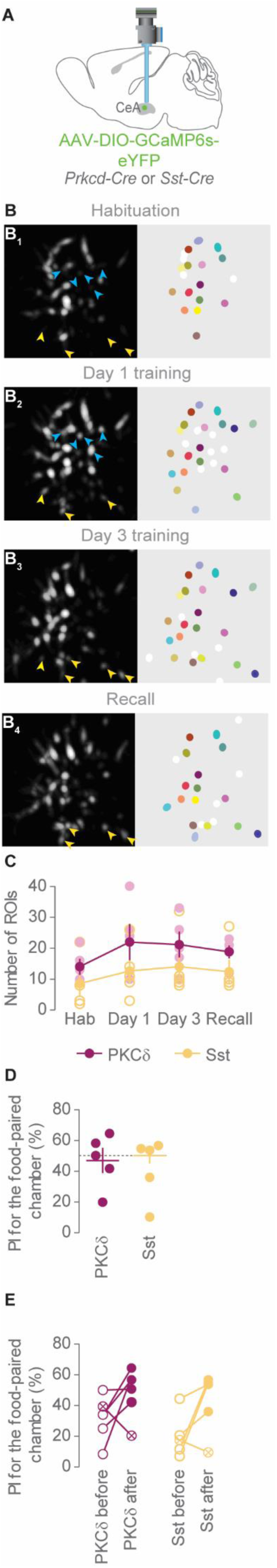
In vivo calcium imaging of CeA^PKCδ^ and CeA^SST^ neurons. (A) Calcium imaging of CeA^PKCδ^ and CeA^Sst^ neurons via miniscope in freely moving mice. (B) Maximum-projection images of PKCδ+ GCaMP6s-expressing neurons from a representative recorded mouse during habituation (B_1_), first (B_2_) and third (B_3_) days of training, as well as recall (B_4_). Corresponding region-of-interests (ROIs) are depicted on the right. ROIs identified over consecutive sessions are shown in identical color. ROIs detected only in one session are shown in white. Blue arrowheads indicate neurons that appeared for the first time on day 1 of conditioning. Yellow arrowheads indicate neurons that are common between at least 2 out of 3 days of conditioning and recall and are not visible on habituation. (C) Numbers of detected ROIs during all four sessions (One-way repeated measures ANOVA, for PKCδ group comparisons: time point, F(3,4) = 2.16, P=0.1458 and for Sst group comparisons: time point, F(3,4) = 3.98, P=0.0351). (D) PI for the context+ during recall for both PKCδ- and Sst-GCaMP6s recorded mice. (E) PI for the context+ before and after conditioning. Data points shown as crossed circles represent mice that did not show an increase in learning after conditioning.(n = 5 PKCδ-GCaMP6s and n = 5 Sst-GCaMP6s recorded animals). Bar graphs show mean +/- s.e.m and each dot is the quantification of a single animal.

We recorded the activities of 24–61 cells per mouse, 202 cells total in 5 Prkcd-cre and 14–60 cells per mouse, 149 cells total in 5 Sst-cre animals (Figure 2C). The numbers of active cells during training and recall were higher compared to habituation day in both groups, probably due to the fact that investigation and consumption of food recruited new sets of neurons (Figure 2B,C). Using a ROIs alignment method (Sheintuch et al., 2017), we registered cell identities across imaging sessions. There were about 10 and 9 % of cells in common for all four recording days in *Pkcd-cre* and *Sst-Cr*e animals, respectively (Figure S2C-D). When excluding the habituation day, the number of common cells for the three recording days increased to 17.3 and 17%, respectively (Figure S2C-D), suggesting that neurons recorded during conditioning and recall may be functionally more similar than the ones initially active during habituation. Of 10 recorded mice, 3 per genotype met the learning criteria, meaning that their preference index for the positive context was 50% or higher during recall (Figure 2D). Nonetheless, 8 animals out of 10 increased their preference for the positive context after conditioning (Figure 2E) indicating a certain degree of learning. It is possible that the weight of the miniscope rendered movements in the arena more laborious and decreased learning performance.

### Central amygdala encoding of Pavlovian appetitive learning

We tested the Hebbian model of appetitive conditioning (Brown et al., 1990; D. O. Hebb, 1949; Sejnowski, 1999) by investigating whether the positive context which represents multisensory information, could serve as a general predictor of food availability after learning. Or, in other words, could CeA neurons that fire during food intake acquire a specific response to the positive context after learning?

Considering only the data of the 8 mice that learned the task (Figure 2E), we convolved the positive context signal with the kinetics of the calcium indicator to create a regressor and correlated across time the activity of each CeA cell with the corresponding regressor (Miri et al., 2011). An example trace of a positive context regressor is shown at the bottom of figure 3L. We found that during habituation, the activities of all neurons showed little correlation to the positive context (Figure 3A-B). Instead, during recall, the activities of both populations shifted toward positive correlations (Figure 3A-B) suggesting that the representations of the positive chamber by CeA^PKCδ^ and CeA^Sst^ neurons were transformed after learning.

**Figure 3:**
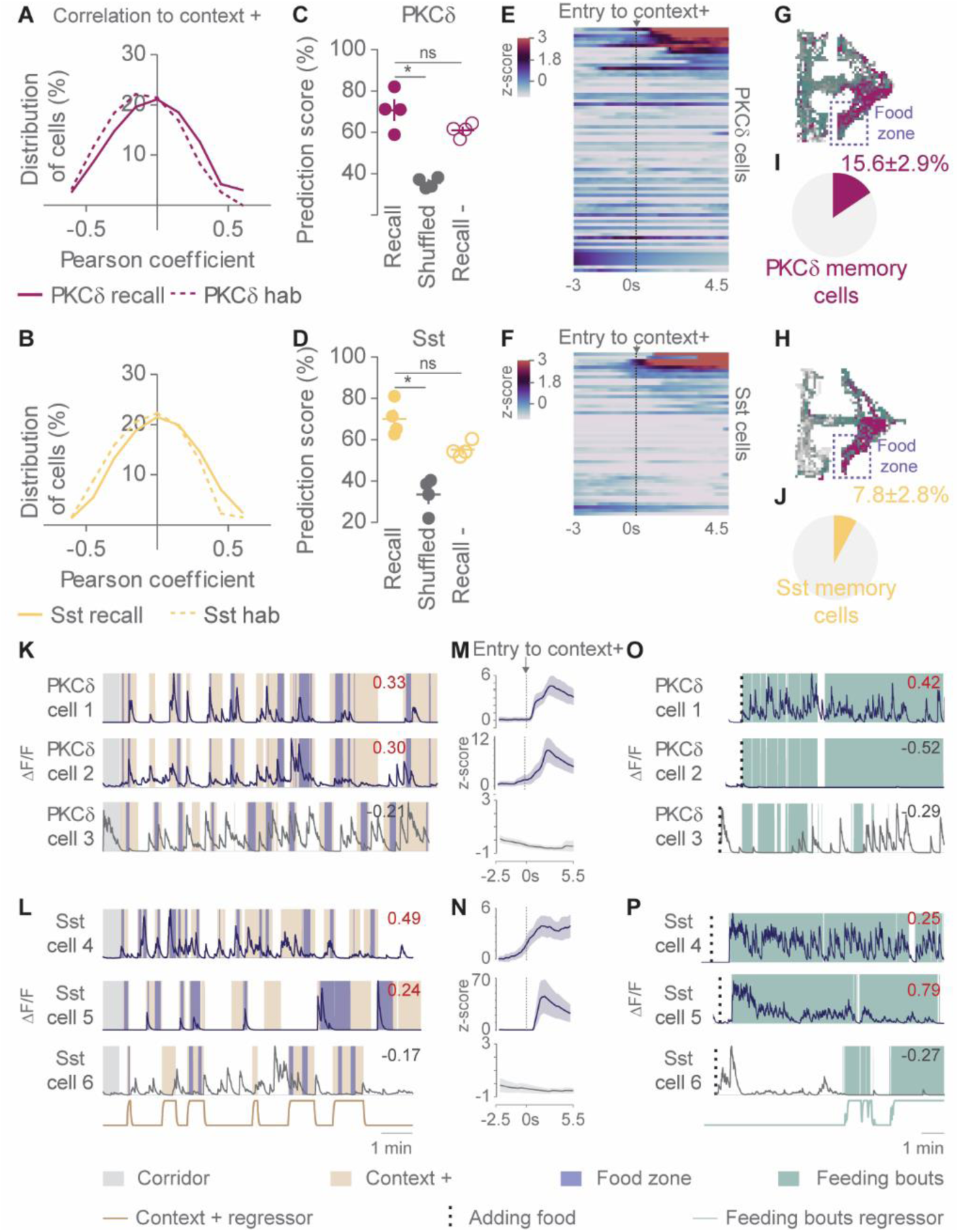
Central amygdala encoding of Pavlovian appetitive learning. (A-B) Frequency distribution of Pearson correlations to the positive context regressor in PKCδ (A) and SST (B) calcium recorded neurons during habituation and recall. (C-D) Prediction scores of the locations of Prkcd-Cre (C) and Sst-Cre (D) calcium recorded animals in the positive context during recall, after temporally shuffling of behavioral data (shuffled) and after excluding 4 neurons per animal that exhibited the highest correlation value to the positive context regressor (recall -) (Friedman test, for PKCδ group comparisons: F= 9.9, P=0.0062 with Dunn’s Multiple Comparison Test and for Sst group comparisons: F = 9.3, P = 0.0115 with Dunn’s Multiple Comparison Test, *P < 0.05). Bar graphs show mean +/- s.e.m and each dot is the quantification of a single animal. (E-F) Heatmap of averaged z-scored calcium responses of PKCδ (E) and SST (F) recorded neurons following entry to the positive context (at 0 sec). Cells were sorted in a descending order based on their activity response upon entry in the context+ (Entries to the context+ varied from 6 to 14 times depending on the mice) (n = 75 PKCδ+ and 50 Sst+ neurons) (G-H) Heat maps of the ΔF/F signal across the whole arena for one representative memory PKCδ (G) and Sst (H) cell. Green represents the minimum and purple the maximum per-pixel frequency. (I-J) Fraction of CeA memory neurons in Prkcd-cre (I) and Sst-cre (J) recorded animals during recall. (K-L) Representative Ca^2+^ traces of single PKCδ (K) and Sst (L) recorded neurons during recall. The trace in beige at the bottom of L represents the convolved GCaMP6s regressor for the positive context of cell 6. (M-N) Average traces of the corresponding PKCδ (M) and Sst (N) neurons shown on the left following entry to the positive context (at 0 sec). Shaded areas represent s.e.m. (O-P) Ca2+ signals of the corresponding PKCδ (O) and Sst (P) neurons shown in figure K-L upon introduction of a food pellet in the food zone (dotted line) at the end of the place preference assay. The trace in green at the bottom of P represents the convolved GCaMP6s regressor for the feeding bouts of cell 6. Values in K-L and O-P indicate the Pearson correlation coefficients of each cell to its corresponding feeding bouts regressor. Values in red indicate a significant correlation. Each ΔF/F trace was normalized to its maximum value. (n = 4 PKCδ- and 4 Sst-GCaMP6s recorded animals)

For each mouse, we randomly selected 70% of the data during recall to train a logistic regression classifier and asked whether the location of the mice in the remaining dataset, could be accurately predicted based on CeA^PKCδ^ and CeA^Sst^ population activity. In all tested mice, we could reliably decode the position of the animals in the positive chamber (70.7 ±4.7% for CeA^PKCδ^ and 70 ±4.1% for CeA^Sst^ recorded ensembles) while performances significantly dropped when the decoder was trained on temporally shuffled behavioral data (Figure 3C-D and Figure S3A) indicating that correct predictions were well above chance. We also found a decreased performance of 10% and 15%, respectively, when the classifier was tested on CeA^PKCδ^ and CeA^Sst^ population activity that excluded the 4 most important features (defined here as the neurons showing the highest correlations to the positive context) (Figure 3C-D). This suggests that a fraction of both populations encodes essential information.

When we aligned the neuronal Ca^2+^ responses of each cell to the onset of the transitions from the neutral to the positive context and averaged the data, we found that a subset of both CeA^PKCδ^ and CeA^Sst^ neurons was activated upon transition to the positive context (Figure 3E-F). These cells exhibited an area-biased activity that was specifically high when the mouse was in the appetitive context and preferentially within the food corner (Figure 3G-H). Cells that displayed a significant correlation toward either the positive context or the food zone regressor or both (memory neurons) accounted for about 16% of CeA^PKCδ^ and 8% of CeA^Sst^ recorded cells during recall (Figure 3I-J). We did not find a group of neurons whose activity increased upon transition to the neutral context (Figure S3B-C).

Analysis of the ΔF/F traces of each memory neuron revealed different patterns of activity within the positive context. Some neurons fired both upon entry to the positive chamber and investigation of the food zone (Figure 3K,M, cells 1 and 2, Figure 3L,N, cell 4). Some cells were active preferentially within the food zone (Figure 3L,N, cell 5). Other neurons showed no preference for the positive context nor the food zone (Figure 3K-N, cells 3 and 6). At this point, no differences were observed between PKCδ and Sst memory cells. Using again a regressor-based approach, we found that the activities of 7/11 of CeA^PKCδ^ and 4/4 of CeA^Sst^ memory neurons were significantly upregulated during food consumption on the same recall day (Fig 3O-P) demonstrating that a subset of these memory cells can be tagged as food responsive.

Memory neurons whose activity represented best the food zone were found in mice that showed lower learning indices, demonstrated as a negative correlation between the average correlation of memory neurons to the food zone regressor and the PI of the mice (Figure S3D). This may reflect that “poor learners” associated only the precise location of the food zone with a rewarding outcome, whereas “good learners” associated the whole positive context or even the arena with food delivery. When we compared the prediction scores of our logistic regression decoder with the PI of the mice, we found that decoding accuracy of the classifier decreased with learning performances (Figure S3E). It is therefore possible that mice that learned best generalized the representation of the positive context to the whole arena.

These results suggest that contextual information paired to the positive context became predictive of food delivery after learning and support a model in which groups of CeA^PKCδ^ and CeA^Sst^ neurons encode contextual food cues in line with an Hebbian’s plasticity mechanism.

### Differences in calcium activity patterns between CeA^PKCδ^ and CeA^SST^ neurons

Next, we examined the activity profiles of all recorded neurons during conditioning, taking into consideration all 5 Prkcd-cre and 5 Sst-cre recorded animals (Figure 2C). On day 1 and day 3 of training, we found that a large fraction of both CeA^PKCδ^ and CeA^Sst^ neurons strongly increased their activity specifically when the mice were sequestered in the positive context where the food reward was present (Figure 4A-B and 4C-D, cells 1 to 6). These cells accounted for 30.9 ±10.3% of PKCδ and 31.1 ± 7.3% of Sst recorded ensembles on day 1 and their proportion increased to 41.7 ± 6.7% and 58.3 ±7.8%, respectively, on day 3 of conditioning (Figure 4E).

**Figure 4:**
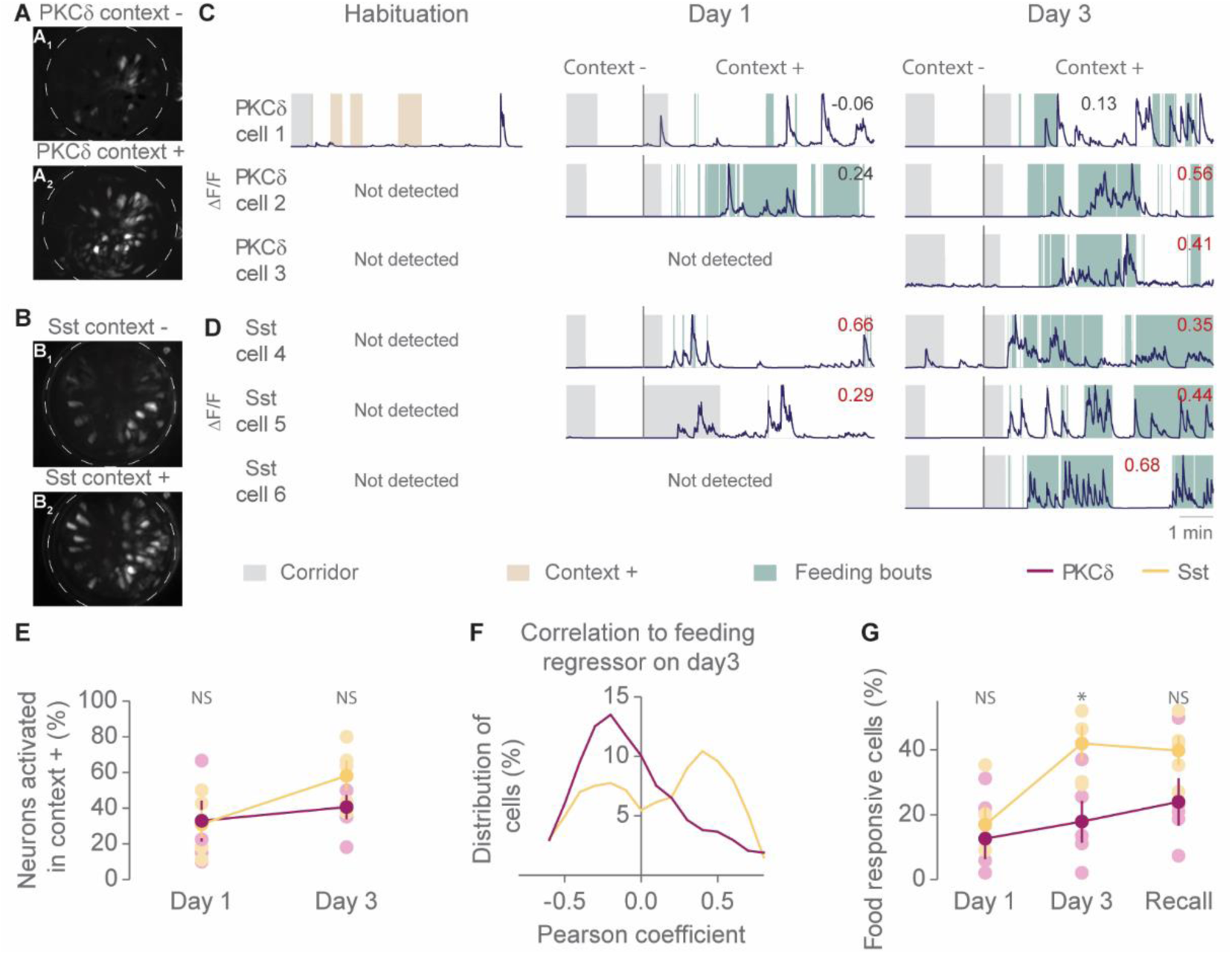
Differences in calcium activity patterns between CeA ^PKCδ^ and CeA^SST^ neurons. (A-B) Maximum-projection representative images of PKCδ+ (A) and Sst+ (B) GCaMP6s-expressing neurons during day 3 of conditioning in the neutral (context -, A_1_ and B_1_) and in the positive context (A_2_ and B_2_). (C-D) Representative Ca^2+^ traces of three PKCδ (C, cells 1-3) and three Sst (D, cells 4-6) recorded neurons during habituation, day 1, and day 3 of conditioning. Note that most cells were not detected during habituation. Colored boxes indicate the location of the mouse in the corridor (grey), in context+ (beige), or on top of the food (green). The values indicate the Pearson correlation coefficients of each cell to its corresponding feeding bouts regressor. Values in red indicate a significant correlation. Each ΔF/F trace was normalized to its maximum value. (E) Proportion of neurons per mouse that are significantly active in context+ compared to context-on days 1 and 3 of conditioning. (Mann-Withney U test between PKCδ and Sst on day1, U = 11, P = 0.8413 and on day3, U = 6, P = 0.2045). (F) Frequency distribution of Pearson correlations to the feeding bouts regressor in PKCδ and SST recorded neurons during day3 of conditioning. (G) Proportion of neurons per mouse that are food responsive on day 1, day 3 of conditioning, and recall. (Mann-Withney U test between PKCδ and Sst on day1, U = 10, P = 0.6723, on day3, U = 2, P = 0.0362 and on recall, U = 4, P = 0.0952). (n = 5 PKCδ- and 5 Sst-GCaMP6s recorded animals). *P < 0.05.

For each cell, we calculated Pearson correlations to their corresponding feeding regressor and classified neurons as activated by food consumption. We found that the majority of cells that showed significant correlation to the feeding regressor on one conditioning day conserved their functional tag on the next day (Figure 4D, cells 4 and 5, and Figure S4A). Most of food responsive neurons were significantly active in the positive context compared to the neutral one during all training sessions and were absent during habituation (Figure 4C, cells 2 and 3, 4D, cells 4 to 6, and Figure S4B).

Interestingly, CeA^PKCδ^ food responsive cells often showed delays in the onset of firing upon food intake and did not fire systematically during each feeding bout (Figure 4C, cells 1 to 3). In comparison, CeA^Sst^ neurons increased their activities at the onset of a feeding bout and more reliably for the next following ones (Figure 4D, cells 4 to 6). Indeed, on day 3 of training, CeA^Sst^ neurons showed stronger positive correlation to the feeding regressor (Figure 4F) compared to CeA^PKCδ^ cells and the proportion of food responsive cells was therefore higher among the CeA^Sst^ population. (Figure 4G) Interestingly there was no difference in the fraction of PKCδ+ and Sst+ cells active in the positive context during training (Figure 4E) suggesting that the activity of CeA^PKCδ^ cells may not directly relate to food consumption but rather to the salience of the food reward. This phenotype was not because Pkcd-Cre animals ate less than Sst-Cre as there was no correlation between the amount of time spent eating and the proportion of food responsive cells on day 3 of training (Figure S4C-D).

When tracing back the origin of the 11 PKCδ and 4 Sst memory cells (Figure 3I-J) we identified two different scenarios:

1. Cells were initially not detected in the arena until day 3 of conditioning where they were specifically active in the positive chamber in the presence of the food pellet. These neurons were tagged as food responsive. They finally developed a preferential activity in the positive context on recall even in the absence of the food reward. This scenario includes 7/11 of the CeA^PKCδ^ and all CeA^Sst^ memory neurons.
2. Cells were initially active or inactive during habituation. During conditioning, they specifically fired in the positive context. These neurons were either not food responsive or lose their “food responsive” tag after conditioning. This scenario includes 4/11 of the PKCδ memory neurons.

These last findings suggest that during conditioning, CeA^Sst^ cells whose activities show high positive correlation with food intake behavior may be important to link environmental information with the physical properties of the food. In contrast, CeA ^PKCδ^ neurons that are robustly activated upon food pellet delivery in the arena, but not reliably upon food consumption, may rather form associations between environmental information and the salience of the food reward.

## DISCUSSION

Here, we used a conditioned place preference assay to study whether neurons in the CeA can encode contextual appetitive memories. We found that two genetically distinct populations, CeA^PKCδ^ and CeA^Sst^ neurons, developed specific activity patterns for the chamber associated with the reward. Interestingly, only the activities of CeA^PKCδ^, but not CeA^Sst^, cells were required for the formation of a preference for the appetitive context, suggesting that these neurons encode a combination of sensory, reward and contextual information to support learning that a spatial location predicts food availability.

These results were rather unexpected, since CeA^PKCδ^ neurons were previously shown to inhibit appetitive behaviors in response to negative events such as threat or aversive tastes, or in response to satiety (Cai et al., 2014; Kim et al., 2017). Moreover, CeA^PKCδ^ neurons could be described as a “broad aversive” population, since these neurons also promote pain-related responses, and are linked to the expression of aversive memories (Haubensak et al., 2010; Wilson et al., 2019). Our findings, instead, suggest that CeA^PKCδ^ neurons may have functions that are more general, relate to a variety of stimuli, and, depending on context, select the appropriate defensive or appetitive responses. Such a model is supported by recent findings on anesthesia neurons in the CeA. Those neurons show a large overlap (nearly 80 percent) with CeA^PKCδ^ neurons and attenuate (rather than promote) pain-related behavior (Hua et al., 2020). CeA^Sst^ neurons are also a good example of a “general encoding” population, since these neurons relate to appetitive stimuli (Kim et al., 2017; Wilson et al., 2019), but also contribute to the generation of defensive behaviors (Li et al., 2013; Penzo et al., 2015; Yu et al., 2016). In the future, further subdivision of CeA populations using two or three genetic markers will be needed to reveal the presence of more specialized cells.

### How could CeA^PKCδ^ neurons mediate learning of contextual food cues?

Inhibition of CeA^PKCδ^ neurons did not affect the numbers of times an animal visited the precise location of the food zone, but rather the time spent within this area. This was consistent with our calcium imaging data showing that CeA memory cells showed a significant increase in activity when the animal was inside the appetitive chamber, either specifically in the food zone or in the positive context. We interpret these results such that the activity of CeA neurons may be more relevant to encode a representation of the goal, i.e. reward associated with a particular location, rather than a representation of compass cues that would help the animal in its search for the food. In the animals that learned best, this memory may initially be linked to the location of the food zone and was then generalized to the positive context and ultimately to the whole arena.

Although contextual memory traces were observed in both CeA^PKCδ^ and CeA^Sst^ neurons, our data uncovered differences between the activity patterns of these two populations. We found, for instance, that the proportion of neurons active during food intake was significantly higher among CeA^Sst^ compared to the CeA^PKCδ^ population. Conversely, the activity of CeA^PKCδ^ neurons often increased somewhat delayed after the onset of feeding possibly at a time when satiety signals start influencing food intake. We hypothesize that CeA^Sst^ cells may be essential to link a context or a sensory stimulus with the physical properties of food (e.g. taste, texture, etc), whereas CeA^PKCδ^ neurons may mediate the association of contextual stimuli with the general affective properties of food, a function that has been previously suggested for the CeA (Balleine and Killcross, 2006). Inhibiting the formation of these memories would then impair an animal’s ability to develop a preference for the location of the food reward.

Of great interest would be now to determine whether the activity of the memory cells is modulated by the animal’s expectation of whether the food pellet will be available and therefore exhibit classic reward prediction error signals (Schultz, 2013). Consistent with this, our calcium data revealed that at least two memory cells reduced their firing upon food consumption during recall suggesting that expectation may suppress responses to food intake after conditioning. In conclusion, our work reports specific activity patterns of CeA neurons that resemble contextual memory traces to appetitive stimuli. Moreover, we refine a model of amygdala function in which “general encoding” subpopulations exist that relate to a variety of stimuli and, depending on context, select the appropriate defensive or appetitive responses.

## ACKNOWLEDGMENTS

We thank Nejc Dolensek, Aljoscha Leonhardt, Matthias Meier and Ruben Portugues for their help with data analysis. We thank J. Cotino and Y. Pignot for their help with management of the animal colony. This study was supported by the Max-Planck Society, the Deutsche Forschungsgemeinschaft (SPP1665), the European Research Council under the European Union’s Horizon 2020 research and innovation programme (No. 885192).

## AUTHOR CONTRIBUTIONS

M.P and F.F designed experiments. M.P and F.F performed experiments. M.P analyzed experiments. R.K supervised experiments. M.P and R.K wrote the manuscript. Funding Acquisition, R.K.

## METHODS

### EXPERIMENTAL MODEL AND SUBJECT DETAILS

#### Animals

Prkcd-Cre (Tg(Prkcd-glc-1/CFP,-Cre)EH124Gsat) BAC mice were imported from the Mutant Mouse Regional Resource Center. Sst-Cre (Ssttm2.1(cre)Zjh) transgenic mice were acquired from the Jackson Laboratory (https://www.jax.org/).Rosa26R44 mouse lines were as described previously (Soriano, 1999). Mice were backcrossed onto a C57BL/6NRj background (Janvier Labs -http://www.janvier-labs.com). 3-6 months old male mice were used for the place preference assay combined with optogenetic manipulation of CeA^PKCδ^ and CeA^Sst^ cells as well as for the open field task. 8 to 18 Months old males and females mice were used for the place preference assay combined with Ca^2+^ imaging of CeA^PKCδ^ and CeA^Sst^ cells. Mice were kept on a 12-h light/dark cycle. All behavior experiments were conducted during the light phase of the cycle and under dimmed light in the behavioral boxes.

#### Viral Constructs

The following AAV viruses were purchased from the University of North Carolina Vector Core (https://www.med.unc.edu/genetherapy/vectorcore: AAV5-ef1a-DIO-eNpHR3.0-mCherry, AAV5-ef1a-DIO-mCherry. The AAV5-Syn.Flex.GCaMP6s virus was obtained from Addgene (http://www.addgene.org/).

## MEDTHODS DETAILS

### Stereotaxic surgeries

#### Viral injections in the CeA

Mice were anaesthetized using isoflurane (Cp-pharma) (induction, 3%; maintenance, 1.5%) in oxygen-enriched air and head-fixed on a stereotaxic frame (Model 1900 – Kopf Instruments). Body temperature was maintained at 37°C using a heating pad. Carprofen (Rimadyl – Zoetis) (5 mg/kg body weight), and an analgesic, were given via subcutaneous injection. Mice were bilaterally injected with 0.3 µl of virus in the CeA by using the following coordinates calculated with respect to the bregma: −1.22 mm anteroposterior, ± 2.9 mm lateral, −4.7 to −4.8 mm ventral. Viral particles were delivered using glass pipettes (708707 - Blaubrand intraMark) connected to a Picospritzer III (Parker Hannifin Corporation) and controlled by a Master-8 pulse stimulator (A.M.P.I) at a flow rate of ∼50 nL/min. After delivery of the virus, the pipette remained in the brain for 5 min to prevent spread of the virus. Virus was allowed to be expressed for a minimum duration of 4.5 weeks and a maximum of 10 weeks before behavioral experiments.

#### Optic fibers implants

Mice used in optogenetic experiments were in the same surgery bilaterally implanted with optic fibers (200-µm core, 0.22 NA, 1.25-mm ferrule - Thorlabs) above the CeA (−4.35 mm ventral from bregma). The skull was first protected with a thin layer of histo glue (Histoacryl, Braun), the fibers were then fixed to the skull using UV light-curable glue (Loctite AA3491 - Henkel) and the exposed skull was covered with dental acrylic (Paladur - Heraeus).

#### GRIN lens implantation and baseplate fixation

For gradient index (GRIN) lenses implantation, 3 weeks after viral injection, mice expressing GCaMP6s in PKCδ+ or SST+ cells were anesthetized using the same procedure as described above. A small craniotomy was made above the CeA using the same coordinates as for the injection of the viral preparation. Debris were removed from the hole and a customized blunted 23G needle (0.7mm in diameter) was slowly lowered down into the brain at a speed of 150 µm/min to a depth of −4.6 mm from bregma. After retraction of the needle, a GRIN lens (ProView lense; diameter, 0.5 mm; length, ∼8.4 mm, Inscopix) mounted on a GRIN lens-holder was slowly (150 µm/min) implanted above the CeA. The skull was first protected with a thin layer of histo glue (Histoacryl, Braun), the lens was then fixed to the skull using UV light-curable glue (Loctite AA3491 - Henkel) and the exposed skull was covered with dental acrylic (Paladur - Heraeus). The exposed top of the lens was protected by a covering of a silicone adhesive (Kwik-cast - World Precision Instruments). 4 to 8 weeks after GRIN lens implantation, mice were anesthetized and placed in the stereotaxic setup. A baseplate (BPL-2; Inscopix) attached to the miniature microscope was positioned above the GRIN lens. Concentration of the anesthetic gas was lowered and the focal plane was adjusted until neuronal structures and GCaMP6s dynamics were clearly observed. Mice were then fully anaesthetized again and the baseplate was fixed using C&B Metabond (Parkell). A baseplate cap (BCP-2, Inscopix) was left in place until imaging experiments.

### Behavioral assays

#### Conditioned place preference

The conditioned place preference behavior was conducted in a custom-built arena made of two-chambers: a rectangle-shaped chamber (45*15 cm) and a triangle shaped chamber (45*30 cm) separated by a corridor (Figure 1A). Chambers additionally differ based on the texture of the floor and pattern on the walls. The surfaces of the two chambers were identical. On the first day of the behavioral experiment, mice were allowed to explore the arena for 20 mins. Preferences for the two chambers were measured and the least preferred one was chosen as the food-paired chamber (context +) for the rest of the paradigm. From then on, and until the end of the experiment, mice were singly housed and food restricted. They were weighed daily and supplied with necessary food to maintain at least 85% of their initial body weight. Conditioning was then conducted over three consecutive days. For this, mice were first sequestered in the neutral context (context -) for 15 mins and then manually transferred to the context + for 15 mins in which they had access to a food pellet of approximately 1 g. The remaining food was weighted at the end of each conditioning session to determine food consumption. On the 5^th^ and last day of the experiment, mice were freely navigating in between the two chambers in the absence of food. The preference for the context + was measured for a period of 10 mins as a readout for contextual appetitive conditioning.

For the optogenetic experiments, PKCδ-eNpH3.0, PKCδ-mcherry, Sst-eNpH3.0 and Sst-mcherry mice received bilaterally, constant 561-nm intracranial light during the whole 30 mins of the conditioning. For this, mice were tethered to optic-fiber patch cables (Doric Lenses or Thorlabs) connected to a 561 nm CNI laser(Cobolt) via a rotary joint (Doric Lenses) and mating sleeve (Thorlabs). During habituation and recall, mice were free of the optic-fiber patch cables.

We excluded 1 PKCδ-eNpH3.0, 1 PKCδ-mcherry and 1 Sst-eNpH3.0 animals that did not at least explore once both chambers during recall.

#### Open-field

PKCδ-eNpH3.0, PKCδ-mcherry, Sst-eNpH3.0 and Sst-mcherry mice were allowed to explore a custom-built plexiglas arena (40 cm × 40 cm × 25 cm) for 10 mins. During the whole experiment, mice received bilaterally, constant 561-nm intracranial light through optic-fiber patch cords (Doric Lenses or Thorlabs) connected to a 561 nm CNI laser(Cobolt) via a rotary joint (Doric Lenses) and mating sleeve (Thorlabs).

### In vivo Ca^2+^ imaging of freely moving mice

All imaging experiments were conducted on freely behaving mice PKCδ-GCaMP6s and Sst-GCaMP6s mice. GCaMP6s fluorescence signals were acquired using a miniature integrated fluorescence microscope system (nVoke - Inscopix). Before each imaging session, the miniscope was secured in the baseplate holder. Mice were habituated to the miniscope attachment procedure 3 days before behavioral experiments. The analog gain and LED output power were adjusted such that the best dynamic fluorescence signals were at the focal plane. Settings were kept constant within subjects and across imaging sessions. Imaging acquisition and behavior were synchronized using the data acquisition box of the nVoke Imaging System (Inscopix), triggered by the Ethovision XT 14 software (Noldus) through a TTL box (Noldus) connected to the USB-IO box from the Ethovision system (Noldus). Compressed images were obtained at 1200 pixels by 800 pixels and 10 frames per second using the Inscopix acquisition software (Inscopix). We recorded the activities of CeA neurons during habituation, day 1 and day 3 of conditioning and recall. To minimize photo-bleaching we limited the duration of acquisition to 3 times 2 mins during habituation, 1 time 2 mins during conditioning in the context - and 3 times 2 mins during conditioning in the context +. Each recording bout was evenly spaced in time. On recall day, Ca^2+^ transients were recorded for the full 20 mins of the behavioral experiment.

### Histology

Animals that underwent behavioral experiments combined with optogenetic manipulations of CeA cells were anesthetized with ketamine/xylazine (Medistar and Serumwerk) (100 mg/kg and 16 mg/kg, respectively) and transcardially perfused with phosphate-buffered saline (PBS), followed by 4% paraformaldehyde (PFA) (1004005, Merck) (w/v) in PBS. Extracted brains were postfixed at 4 °C in 4% PFA (w/v) in PBS for 12 h embedded in 4% agarose (#01280, Biomol) (w/v) in PBS and sliced using a Vibratome (VT1000S - Leica) into 100-μm free-floating coronal sections.

Mice that underwent calcium imaging experiments were anesthetized with ketamine/xylazine (Medistar and Serumwerk) (100 mg/kg and 16 mg/kg, respectively) and decapitated. The head together with GRIN lens implant and baseplate, were fixed at 4 °C in 4% PFA (w/v) in PBS for a minimum of 4 days before dissection of the brain. Extracted brains were sliced using a Vibratome (VT1000S - Leica) into 100-μm free-floating coronal sections.

### Microscopy

Epifluorescence images were obtained with an upright epifluorescence microscope (Zeiss) with 5×/0.15 or 10×/0.3 objectives (Zeiss). Images were minimally processed with ImageJ software (NIH) to adjust for brightness and contrast for optimal representation of the data. A median filter was used to decrease noise.

## QUANTIFICATION AND STATISTICAL ANALYSIS

### Extraction of behavioral data for the conditioned place preference assay

In the conditioned place preference assay, the animal location was recorded at 15Hz using a webcam (Logitech) suspended above the arena and velocity as well as the preference index and number of entries to the positive context and the food zone were automatically analyzed by Ethovision XT 14 software (Noldus).

For analysis of the calcium recordings, we could not directly score feeding bouts since we only used a camera that was suspended above the arena. Therefore, we manually scored the time when the animals were on top of the food and choose a criteria of 2.5s as a minimum time to define a feeding bout.

### Extraction of behavioral data for the open-field task

Animal location was recorded at 15Hz using a webcam (Logitech) suspended above the arena and the number of entries to the center of the arena (20 cm × 20 cm square), cumulative duration in center as well as velocity were assessed with Ethovision XT 14 software (Noldus).

### Extraction of ΔF/F and temporal registration with behavioral data

For imaging data processing and analysis, all videos recorded from one imaging session were combined into a single image stack using the inscopix data processing software (version 1.3.0 – Inscopix) and saved as a tiff. Tiff files were then processed using the miniscope 1-photon imaging signal extraction pipeline (MIN1PIPE) (Lu et al., 2018) which returns fully processed ROI components with spatial footprints and temporal calcium traces as outputs. Briefly, the data go through different steps of neural enhancing, hierarchical movement correction, and neural signal extraction that combine a first seeds-cleansing step followed by a simplified spatiotemporal CNMF. Behavioral data were finally temporally aligned to the calcium traces using linear interpolation and unix time stamps as references for both datasets.

### Longitudinal registration of ROIs

ROIs from several recording sessions were longitudinally registered using CellReg Matlab GUI (Sheintuch et al., 2017) (https://github.com/zivlab/CellReg). In brief, the roifn output variable from the Min1pipe that contains the processed vectorized ROI footprints for each session was transformed in matlab using: roi_use = permute(reshape(roifn, pixh, pixw, n), [3, 1, 2]), where n is the number of ROIs.

Transformed ROIs footprints were then registered using CellReg and the following parameters: alignment type: translations and rotations (max rotation in degrees: 30). A maximal distance of 12 microns was used to compute the probabilistic model. The initial and final cell registrations were performed using spatial correlation models. The resulting cell_to_index_map file was used to identify identical ROIs from one day to another, calculate the total number of recorded cells per animal and the number of overlapping neurons during the whole recording session.

### Regressors and correlation analyses

Regressors were built as previously described (Miri et al., 2011). For this, the behavior of each mouse in the positive context, food zone or “on top of the food” (square-wave data sets) was convolved with a kernel with an exponential decay based on the measured half-decay time for GCaMP6s (∼0.150 s) (Chen et al., 2013). The resulting predicted calcium traces were then used to compute Pearson correlation coefficients with the corresponding calcium traces. To classify neurons as “memory cells” or “food responsive cells”, we examined which coefficients arise above chance by correlating our fluorescent traces to 1000 random regressors that were constructed after randomly shuffling the real behavioral dataset by bouts of 10s. We required a threshold of at least 2.58 deviation from the standard error of the random coefficients mean (corresponding to the 99% confidence interval) to assign a cell to a particular functional group.

### Classification of neurons preferentially active in the positive context

To quantify which neurons were preferentially active in the positive context compared to the neutral one during conditioning, ΔF/F transients were z-scored at each time point using the following formula: (F(t) – Fm)/SD, where F(t) is the ΔF/F value at a time t, Fm, and SD are the mean and standard deviation of the baseline calculated from time point when the animals were in the neutral context. An average of the single z-scored time points was then calculated for when the animal was located in the positive context. Neurons were considered to be preferentially active in the positive context when the averaged z-scored value exceeded the 1.96 threshold (corresponding to the 95% confidence interval).

### Decoding of positive context locations

To decode the location of the mice in the positive context during recall, we used a logistic regression classifier. For decoder training and testing, we used neuronal Ca^2+^ signals expressed as ΔF/F. For each animal, classifiers were trained on 70% of the data during recall and tested on the remaining 30%. We computed the prediction score as the average of correct predictions over a 10-fold cross-validation procedure. Correct predictions were defined as the ratio of TP / (TP + FP +FN) where TP is the number of true positives, FP the number of false positives and FN the number of false negatives.

To evaluate the statistical significance of decoding performance, we trained logistic regression decoders on temporally shuffled behavioral data. For this, behavioral data were split into 7 sec bouts and randomly shuffled. This was repeated 5 times. The shuffled prediction score was defined as the average of correct predictions of these 5 repetitions.

### Alignment of calcium responses to positive context entries

ΔF/F transients were z-scored with the baseline calculated from time points when the animals were in the neutral context (see formula above in: Classification of neurons preferentially active in the positive context). We omitted short bouts whose duration was below 2.5 sec to exclude epoch when the animals were only shortly going in and out of the corridor space without really entering the positive context. Z-scored calcium responses of single neuron to positive context entry were then averaged in a time window from 3 s before transition to 4.5 s after. Cells were finally sorted in a descending order based on their activity response upon entry in the context+.

### Heatmaps of the spatial Ca2+ activity

To plot a heatmap of the average spatial activity of one selected cell we used the raw ΔF/F data. The total activity in a specific x-y location was normalized to the total time the animal spent in that location. x-y data were discretized in 50 x 50 pixels.

### Statistical analysis

No statistical methods were used to pre-determine sample sizes. The numbers of samples in each group were based on those in previously published studies. Behavioral experiments were conducted by an investigator with knowledge of the animal genotype and treatment. For behavioral and *in vivo* imaging experiments, behavioral-tracking software and custom-written Python scripts were used to obtain and analyze the data in an automated and unbiased manner. Statistical analyses were performed with Prism 7 (GraphPad) and all statistics are indicated in the figure legends. t-tests were used for individual comparisons of normally distributed data. When normality was not assumed, Mann-Whitney U test and Wilcoxon signed-rank test (for paired observations) were performed for individual comparisons. A one-way repeated measures ANOVA or Friedman’s test (as a non-parametric equivalent) were used for within-subjects comparisons followed respectively by Bonferroni post hoc analysis or Dunn’s multiple comparison test. After the conclusion of experiments, virus-expression, optic-fiber and GRIN-lens placement were verified. Mice with very low or null virus expression as well as mice in which the optical fibers were wrongly located or at least more than 500 µM above the CeA were excluded from analysis.

## DATA AND CODE AVAILABILITY

The datasets supporting the current study have not been deposited in a public repository but are available from the corresponding author on request. This study used custom-built Python 3.0 programmed scripts that are available from the Lead Contact upon request.

## SUPPLEMENTAL FIGURES AND LEGENDS

**Supplementary Figure 1:**
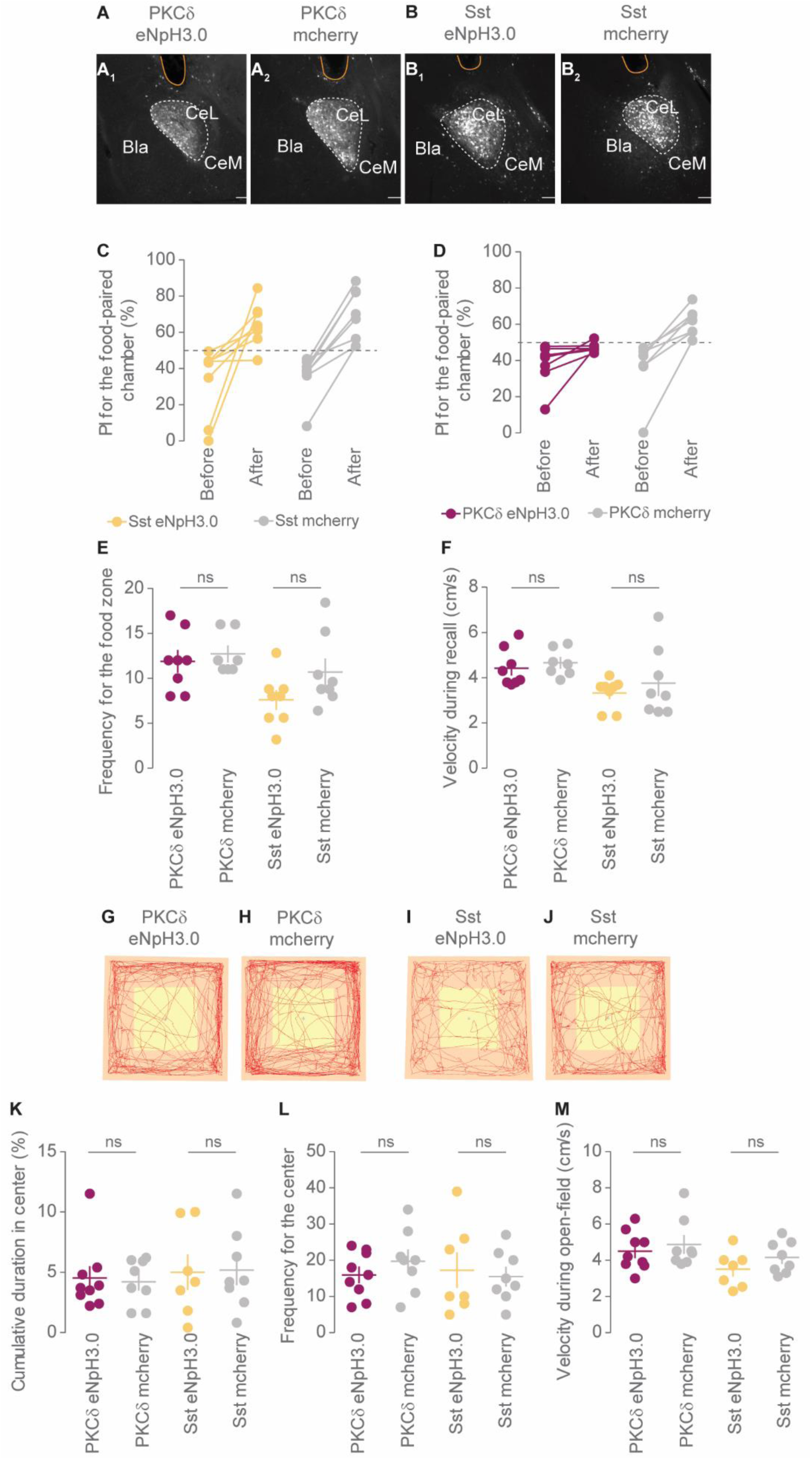
(A-B) Representative epifluorescent images of Prkcd-Cre (A) and Sst-Cre mice (B) injected in the CeL with an AAV-DIO-eNPH3.0-mcherry (A_1_ and B_1_) or AAV-DIO-mcherry (A_2_ and B_2_) and showing the location of the optic fiber track (in yellow). Scale bars: 100 µm. (C-D) PI for the context+ before and after conditioning for CeA^Sst^ (C) and CeA^PKCδ^ neuron-inhibited animals (D), and respective control mice. (E) Number of visits in the food zone during recall (two-tailed unpaired t-test, for PKCδ group comparison: t(13) = 0.5618, P = 0.5838 and for Sst group comparison: t(14) =1.772, P = 0.0982). (F) Velocity during recall (two-tailed unpaired t-test, for PKCδ group comparison: t(13) = 0.6142, P = 0.5497 and for Sst group comparison: t(14) =0.7520, P = 0.4645). (G-J) Representative traces (in red) of the behavior of PKCδ-eNpH3.0 (G), PKCδ-mcherry (H), Sst-eNpH3.0 (I) and Sst-mcherry mice (J) during the openfield task. The yellow square represents the center. (K) Cumulative duration in center during the openfield task (Mann-Withney U test, for PKCδ group comparison: U = 33, P = 0.8096 and for Sst group comparison: U = 25.5, P = 0.8168). (L) Number of visits in the center during the openfield task (two-tailed unpaired t-test, for PKCδ group comparison: t(15) = 1.033, P = 0.3181 and for Sst group comparison: t(13) =0.3481, P = 0.7334). (M) Velocity during the openfield task (two-tailed unpaired t-test, for PKCδ group comparison: t(15) = 0.6166, P = 0.5467 and for Sst group comparison: t(13) =1.348, P = 0.2006). (For the place preference assay, n = 8 PKCδ-eNpH3.0 and 7 PKCδ-mcherry mice and n = 8 Sst-eNpH3.0 and n = 8 Sst-mcherry mice) (For the open-field task, n = 9 PKCδ-eNpH3.0 and 8 PKCδ mcherry mice and n = 7 Sst-eNpH3.0 and n = 8 Sst-mcherry mice). Bar graphs show mean +/- s.e.m and each dot is the quantification of a single animal.

**Supplementary Figure 2:**
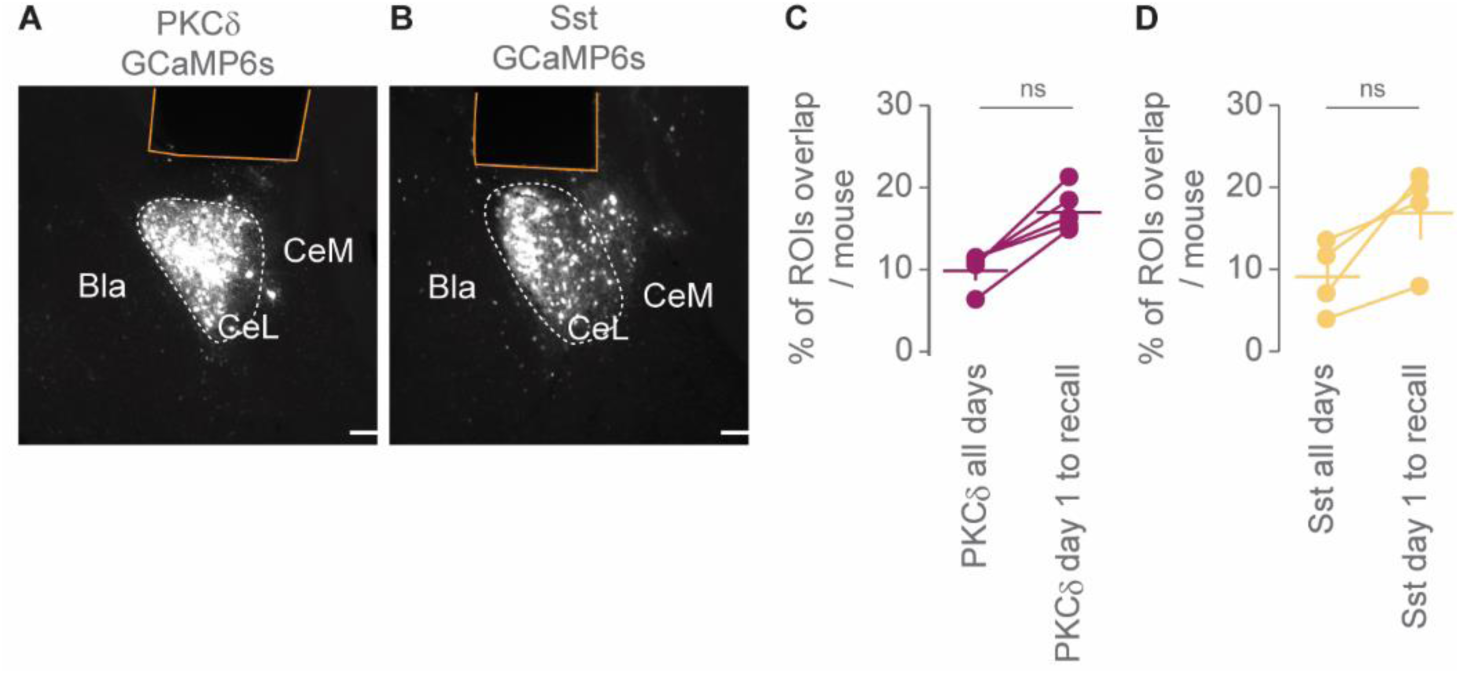
(A-B) Representative epifluorescent images of Prkcd-Cre (A) and Sst-Cre mice (B) injected in the CeL with an AAV-DIO-GCamp6s-eYFP and showing the location of the GRIN lens track (in yellow). Scale bars: 100 µm. (C-D) Percentage of ROIs per Prkcd-Cre (C) and Sst-Cre (D) mouse that overlapped in all four recording sessions or from day 1 of conditioning to recall (Wilcoxon signed-rank test, for PKCδ comparison: P=0.125, and for Sst group comparison: P = 0.25). (n = 5 PKCδ- and 5 Sst-GCaMP6s recorded animals). Bar graphs show mean and each dot is the quantification of a single animal.

**Supplementary Figure 3:**
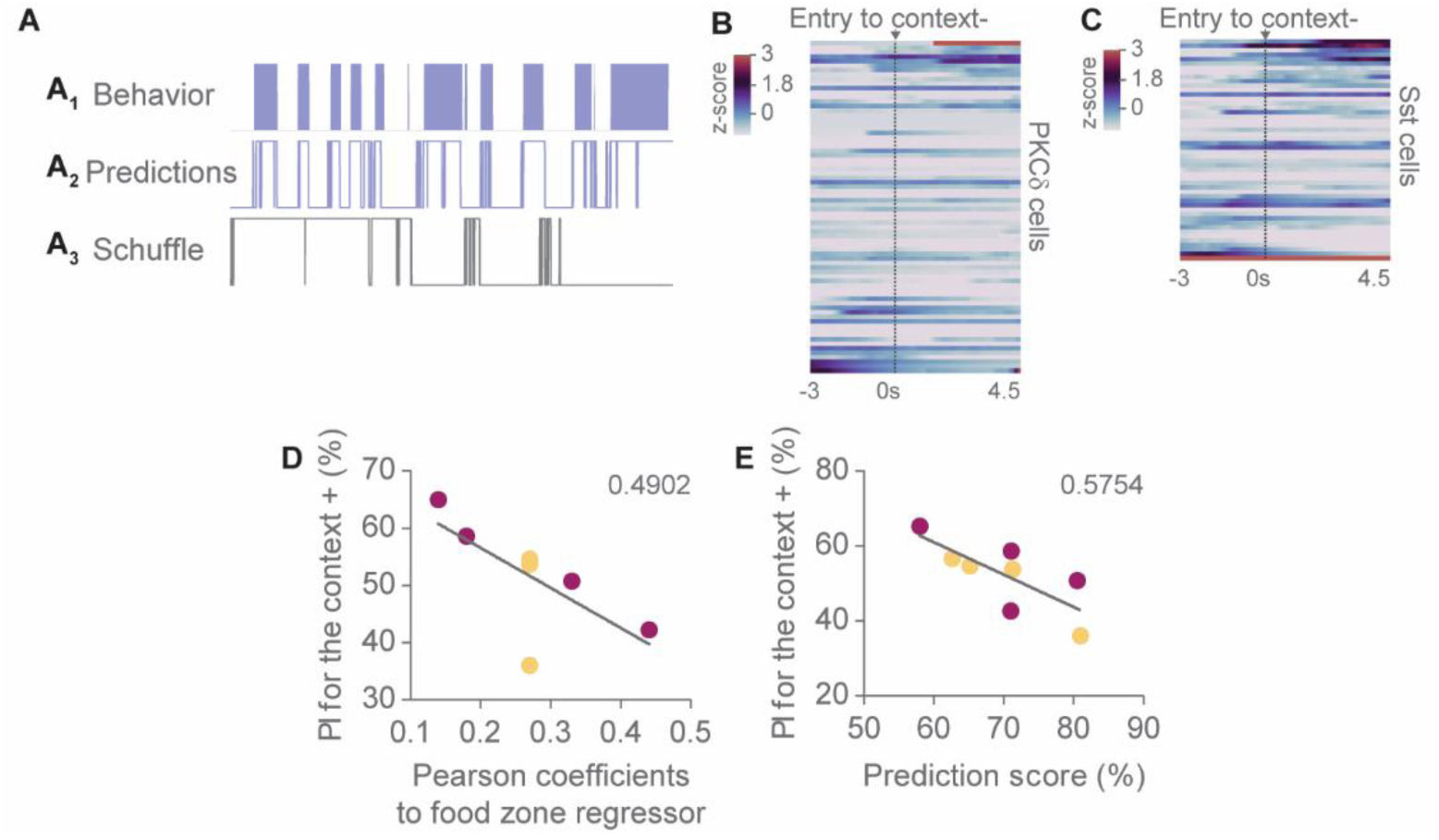
(A) Traces showing the behavior of one representative animal in the positive context during recall (A1), the corresponding predictions by the logistic regression classifier (A2) and the corresponding predictions after randomly shuffling the behavior data (A3). (B-C) Heatmap of averaged z-scored calcium responses of PKCδ (B) and SST (C) recorded neurons following entry to the neutral context (context- at 0 sec). ΔF/F transients were z-scored with the baseline calculated from time points when the animals were in the positive context. Cells were sorted in a descending order based on their activity response upon entry in the context-(n = 75 PKCδ+ and 50 Sst+ neurons). (D) PI on recall day in function of Pearson correlation coefficient value to the food zone regressor. Each dot is the quantification of a single Prkcd-Cre (purple) or Sst-Cre (yellow) animal. Values shown are R square. (E) Prediction score of the logistic regression classifier in function of the PI of the animal on recall day. Each dot is the quantification of a single Prkcd-Cre (purple) or Sst-Cre (yellow) animal. Values shown are R square. *P < 0.05.

**Supplementary Figure 4:**
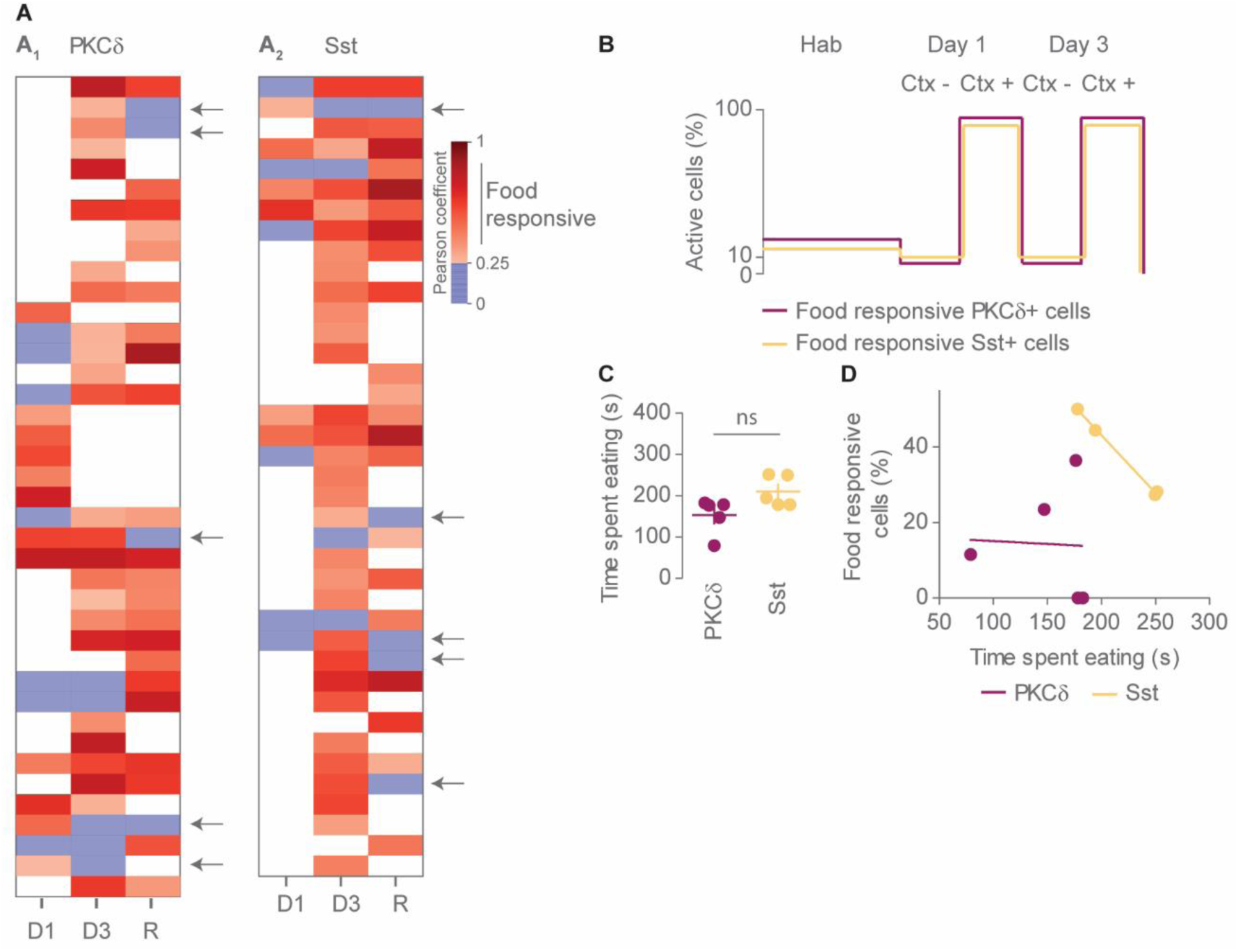
(A) Heatmap of Pearson correlation coefficients of PKCδ (A_1_) and SST (A_2_) temporally registered neurons to the feeding bouts regressor. Blue represents not significant correlations so non-food responsive cells. Strong correlation values are represented in dark red. Arrows indicate neurons that were classified as food responsive on one day and lost their functional tag on the following one. (n = 40 PKCδ+ and 39 SST+ neurons) (B) Proportion of food responsive PKCδ+ and SST+ neurons that are active during habituation, day 1 and day 3 of conditioning and in the context- or the context+ (n = 40 PKCδ+ and 39 SST+ neurons). (C) Time spent eating during day 3 for PKCδ- and Sst-GCaMP6s recorded mice (two-tailed unpaired t-test t(8) =2.242, P = 0.552). Bar graphs show mean +/- s.e.m and each dot is the quantification of a single animal. (D) Proportion of food responsive PKCδ+ and SST+ cells in function of the time spent eating on day 3 of conditioning. Each dot is the quantification of a single animal. (n = 5 PKCδ- and 5 Sst-GCaMP6s recorded animals).

